# LENS: a transferable neuromuscular signature of musculoskeletal pain intensity

**DOI:** 10.64898/2026.09.08.750203

**Authors:** Pedram Mouseli, Nathaniel W. De Vera, Suha Sagheer, W. Darlene Reid, Igor Jurisica, Martin S. Angst, Nima Aghaeepour, Massieh Moayedi, Iacopo Cioffi

## Abstract

Self-reported pain remains the clinical standard for assessing musculoskeletal pain severity, but its subjectivity limits objective diagnosis, monitoring, and therapeutic development. We present LENS (Latent sEMG Neuromuscular Signature), which decodes exertion-evoked musculoskeletal pain intensity from surface electromyography (sEMG) alone. LENS was pre-trained via a cross-modal self-supervised objective that reconstructs muscle oxygenation from sEMG in 183 adults. LENS was trained exclusively on healthy muscle physiology and generalized to an independent healthy cohort (Spearman’s ρ=0.52) and discriminated moderate-to-severe pain (AUROC=0.83). Using a label-free test-time adaptation, LENS generalized to an unseen chronic musculoskeletal pain cohort—myogenous temporomandibular disorder (mTMD)—with comparable performance (ρ=0.61; AUROC=0.86). Reverse transfer, from mTMD to controls, was substantially weaker, an asymmetry attributable to elevated motor variability in chronic pain; excluding highly variable mTMD participants recovered performance, indicating a conserved pain signature masked by pathology-specific motor noise. LENS offers a scalable, physiological correlate of musculoskeletal pain from a single sEMG channel.

---

Musculoskeletal (MSK) pain disorders are among the leading causes of years lived with disability worldwide^1^. Temporomandibular disorders (TMDs), as the second most common chronic MSK pain condition^2^, degrade quality of life through masticatory muscle pain (myalgia) and dysfunction. Like other MSK pain conditions, assessment of TMDs rests almost entirely on self-report. Pain is defined as a subjective experience^3^, and self-report is therefore indispensable^4^. However, it is constrained by inter-individual differences in how sensation is scaled^5^, by cognitive and contextual reporting biases^6–8^. This underscores the need for objective physiological measures that can complement, rather than replace, self-reported pain^9–11^. In myogenic pain, peripheral physiological signals captured from muscle offer a potentially accessible and scalable avenue toward this goal.

Surface electromyography (sEMG) and near-infrared spectroscopy (NIRS) non-invasively capture neuromuscular and hemodynamic changes that accompany exertion, fatigue and pain^12–19^. Previous work has shown that painful TMDs are accompanied by altered motor patterns and abnormal muscle hemodynamics^20–24^. However, translating these observations into reliable predictions for individual people has been challenging^25,26^. Machine-learning studies in this domain reduce continuous recordings to hand-crafted summary statistics, which fail to fully exploit the temporal structure of the signal and its coupling to metabolic state^19,27–29^. These studies have also been confined to small, homogeneous cohorts and single hardware configurations, so transferability to new individuals, cohorts, pathologies or devices remains untested. Validated brain-based signatures of pain now exist^30–33^, but no peripheral counterpart has been established.

Whether peripheral muscle activity contains a pain representation conserved across health and chronic pain remains unknown. Chronic MSK pain involves central mechanisms^34^, including central sensitization^35^, that can amplify peripheral nociceptive input from muscles, resulting in a disconnect with patient reports. Pain also results in protective (i.e., nocifensive) behaviours and altered recruitment strategies^36,37^, which can influence the recorded signal. Therefore, a generalizable peripheral signature could fail in two ways: the encoding could either be altered in patients, or preserved but obscured by pathology-specific motor variability. A stringent test of such a signature is therefore whether a model trained exclusively on healthy muscle can generalize to chronic pain without retraining.

Our modelling strategy was designed around three challenges to achieving this generalization. First, perceived MSK pain is cumulative and depends on the history of exertion rather than the instantaneous state of the muscle. It must therefore be learned from the raw time–frequency signal, with long-range temporal dependencies represented^38–41^. Second, electrical and hemodynamic responses reflect distinct but physiologically coupled aspects of muscle function^42–44^. We therefore used simultaneously recorded NIRS as a cross-modal self-supervised learning target, encouraging the sEMG encoder to learn representations that reflect this shared physiology while requiring only sEMG at deployment. Third, peripheral signals vary substantially across individuals because of anatomical and physiological differences. Supervised models often rely on labelled target-domain data to compensate, whereas we use a brief, label-free test-time adaptation step.

Here, we present LENS (Latent sEMG Neuromuscular Signature), a peripheral neuromuscular signature of MSK pain intensity. LENS was developed from synchronized sEMG and NIRS recordings acquired during a standardized jaw-clenching task, using muscle oxygenation only during pre-training. A model trained exclusively on healthy participants’ pain labels predicts pain in individuals with chronic mTMD which supports the existence of a conserved peripheral representation of MSK pain intensity. LENS also tracks self-reported pain in held-out healthy adults and outperforms classical feature-engineering baselines. However, reverse transfer from chronic mTMD to healthy individuals fails. We attribute this directional asymmetry to elevated within-participant motor variability in mTMD. Model attributions localize the learned representation to high-frequency (150–400 Hz) motor activity, whose decline across repeated exertion scales with reported pain intensity. Together, these findings identify a conserved peripheral representation of MSK pain that can be learned from a single sEMG channel per muscle.

## Results

### A multimodal dataset and a cross-modal learning strategy

To decode the physiological correlates of subjective pain, we created a multimodal dataset of experimentally induced orofacial pain, comprising approximately 72.5 hours of synchronized sEMG and NIRS recordings. Data were collected from the masseter muscles of 183 participants, divided into a training cohort of 81 healthy controls (Control-train), a distinct pathological cohort of 54 individuals with chronic mTMD, and a hold-out test cohort of 48 healthy controls (Control-test) (Fig. 1a). Control-train and mTMD data came from a larger study that also included quantitative sensory testing and brain imaging, not analysed here. Control-test was collected independently. The mTMD cohort tests whether the signature is conserved or obscured by chronic pathology.

**Fig. 1:**
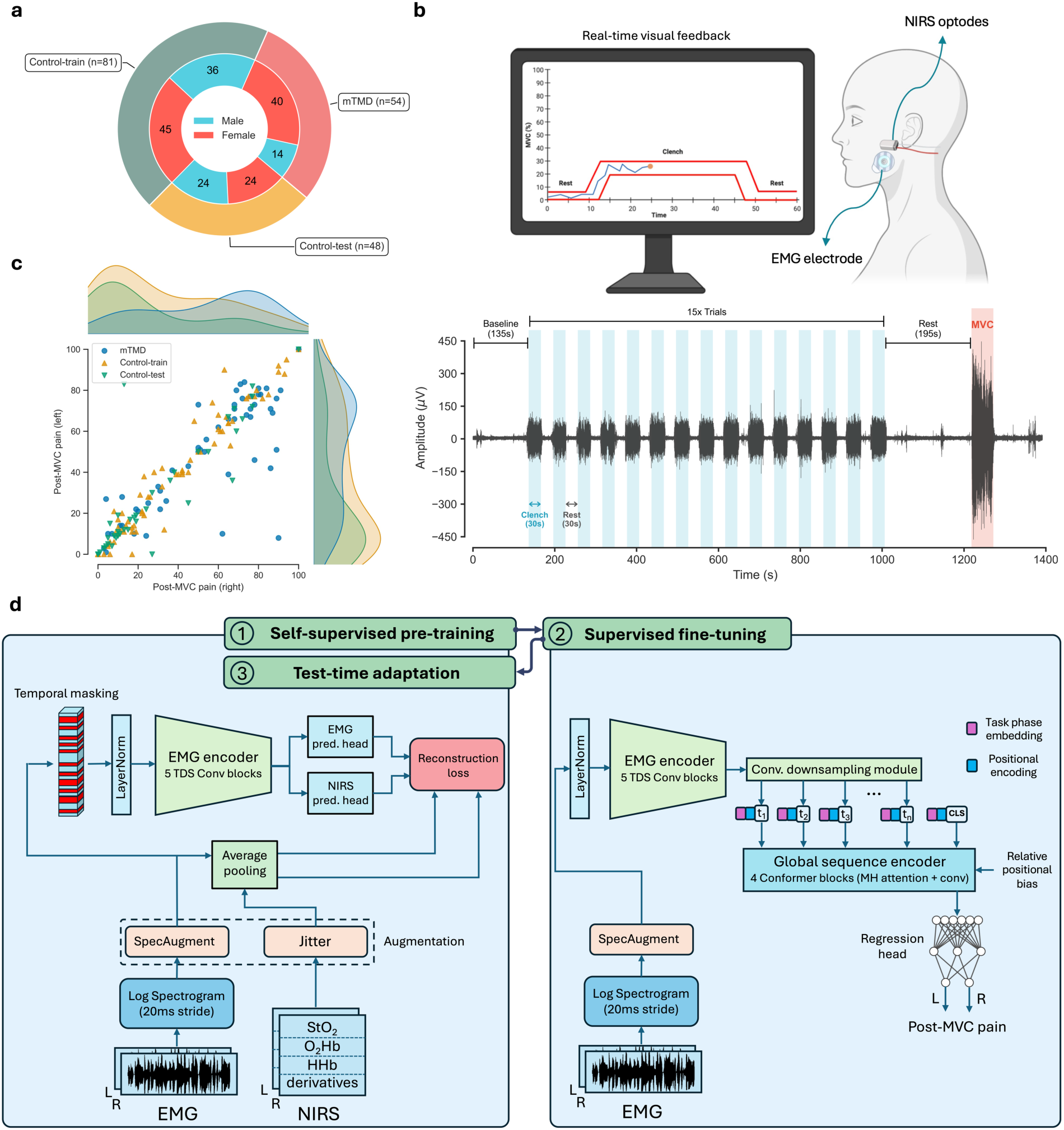
Experimental protocol and hierarchical deep learning architecture. **a.** Cohort composition showing sample sizes for mTMD (n=54) and healthy controls (Train n=81, Test n=48). **b.** Data acquisition. Participants (n=183) performed a repetitive sub-maximal tooth clenching task (15 trials: 30s contraction, 30s inter-trial rest) followed by a sustained maximum voluntary contraction (MVC) until failure. Synchronized sEMG and NIRS were recorded from the masseter on both sides; subjective pain (0–100 VAS) was reported during inter-trial rest periods and after the MVC. **c.** Distribution of post-MVC pain ratings, illustrating a heavy-tailed, non-unimodal profile. **d.** Model architecture. A time-depth separable (TDS)^46^ convolutional encoder extracts local temporal features, which are temporally downsampled and then processed by a Conformer^39^ (Convolution-augmented Transformer) to model long-range dependencies. Training occurs in three stages: (1) Cross-modal self-supervised pre-training (reconstructing EMG and NIRS from masked EMG), (2) Supervised fine-tuning for pain regression, and (3) test-time adaptation (TTA) to individual physiology. Stage 3 reuses the Stage 1 objective and is therefore drawn within the same panel. The NIRS branch is optional for Stage 3. Adaptation using the sEMG reconstruction term alone gives equivalent performance on the primary endpoint (Table S1).

The experimental paradigm was designed to induce a progressive, cumulative pain experience driven by repetitive exertion and muscle fatigue. Participants performed a standardized low-level repetitive jaw clenching task (15 trials of 30s contraction, 30s rest), followed by a sustained jaw clenching at maximum voluntary contraction (MVC) until exhaustion (Fig. 1b). Real-time visual feedback of EMG amplitude was derived from the right masseter alone, referenced to a right-side MVC calibration, and participants were not told which side drove the display. Signals were recorded bilaterally and pain was rated separately for each side, so the right masseter operated within a closed feedback loop while the left provided a free signal under the same protocol. We therefore designate the left masseter as the primary endpoint and report the right as secondary. Pain ratings (100-mm visual analog scales; VAS^45^) were collected after each clench trial (during rest periods) and after the MVC. The resulting distribution of pain labels was highly skewed, characterized by a large concentration of near-zero ratings and a broad spread of higher pain scores in controls and the opposite in mTMD (Fig. 1c). Most sEMG decoding research relies on high-density arrays of tens to hundreds of electrodes; all recordings here used a single concentric or bipolar surface electrode per masseter—i.e., two channels in total—acquired with two distinct hardware systems (Device A and Device B; see *Methods*), testing robustness to hardware-driven artefacts.

To map these signals to subjective pain ratings, we developed a hierarchical deep learning framework (Fig. 1d). Log-spectrograms of the sEMG signal pass through a time-depth separable convolutional encoder^46^ and then a conformer^39^, which weighs task phases—rest, low-level clenching and maximal exertion—across the experimental timeline, so that the cumulative history of exertion rather than the instantaneous state of the muscle informs the prediction.

Training proceeds in three stages (Fig. 1d). In cross-modal self-supervised pre-training, the encoder reconstructs masked segments of both the EMG signal and the simultaneous NIRS hemodynamics (oxygenated and deoxygenated hemoglobin, tissue oxygen saturation, and their rates of change). This objective embeds neurovascular coupling—the relationship between electrical neural drive and metabolic oxygen consumption—into the EMG encoder, and because that context is internalized during pre-training, NIRS is not required for supervised fine-tuning or for inference. Supervised fine-tuning then trains the encoder and conformer end to end to predict post-MVC pain intensity, and test-time adaptation updates the encoder on each test participant’s unlabelled data. Test-time adaptation reuses the pre-training objective and can be run with or without its NIRS term, with equivalent performance on the primary endpoint (Table S1).

### LENS decodes exertion-evoked pain and outperforms classical baselines

We first evaluated the model’s ability to decode subjective MSK pain intensity within specific cohorts. The model, trained on the healthy “Control-train” dataset and evaluated on the independent “Control-test” dataset, successfully predicted post-MVC pain intensity (left: ρ=0.52, p < 0.001; right: ρ=0.48, p < 0.001; Table 1). Across all six evaluation settings, predictive performance on the free left masseter equalled or exceeded that on the feedback-entrained right masseter (Table 1). The opposite pattern would be expected if the model were exploiting feedback-driven regularities, supporting the left masseter as the primary endpoint. Importantly, this predictive capability maintained robust performance across both devices (Fig. 2a), showing generalization across a test set with a markedly different distribution of recording hardware (Fig. 2b). This indicates that the learned features are physiological rather than hardware artefacts.

**Fig. 2:**
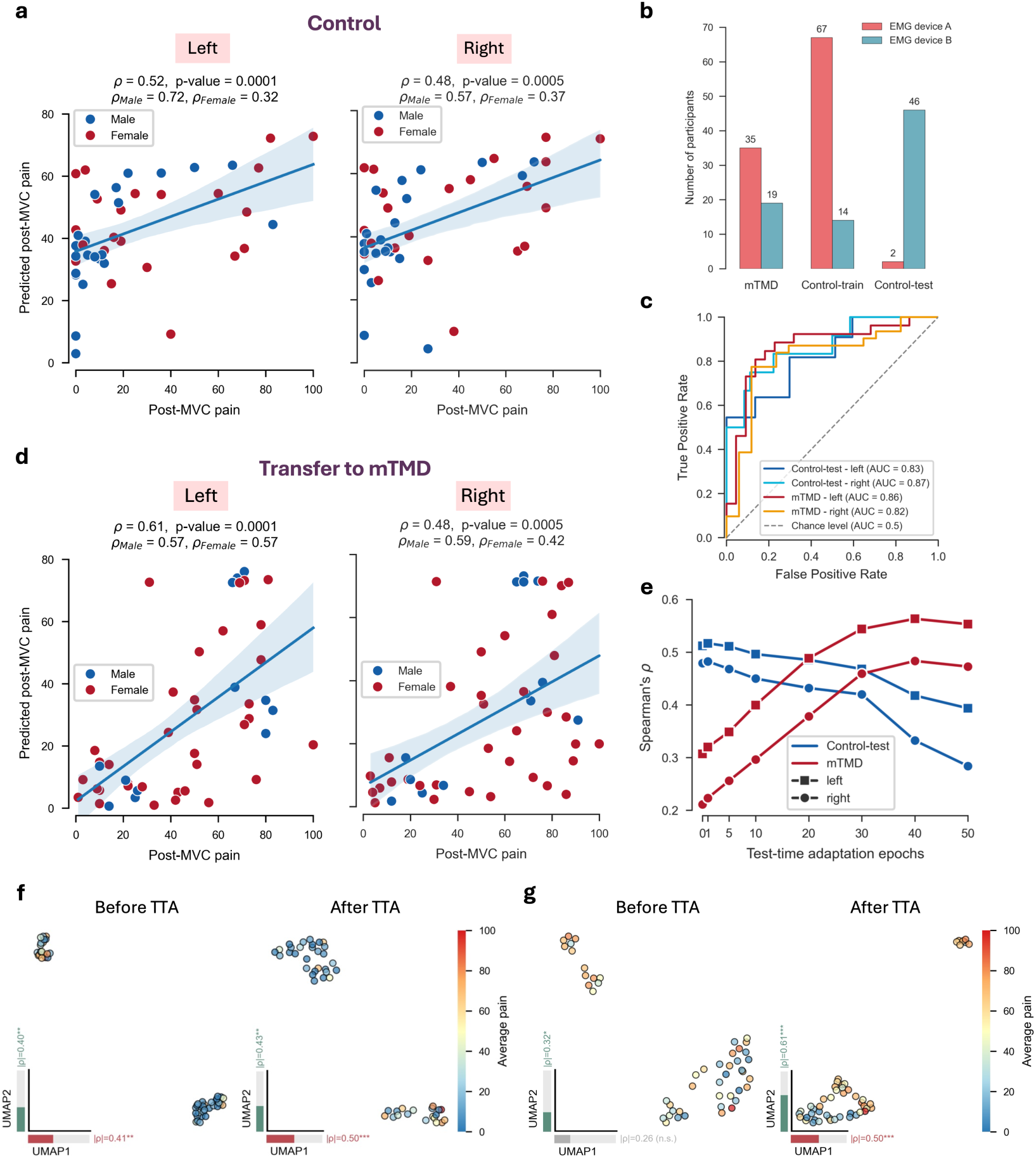
Dataset heterogeneity and cross-domain pain prediction. **a.** Model performance on the healthy hold-out set (Control-test). Each point represents one participant; the model robustly predicts pain (*_ρ_* ≈ 0.5). **b.** Distribution of recording hardware (Device A vs. Device B), highlighting technical heterogeneity. **c.** Receiver Operating Characteristic (ROC) curves evaluating the model’s ability to classify moderate-to-severe pain (VAS≥45/100). The model exhibits strong diagnostic discrimination in both the healthy hold-out (Control-test; left side AUC = 0.83, right side AUC = 0.87) and pathological transfer (mTMD; left side AUC = 0.86, right side AUC = 0.82) cohorts. **d.** Cross-domain generalization. The model trained only on healthy controls effectively predicts pain in unseen mTMD participants, substantially outperforming the reverse transfer (mTMD to Control). **e.** Adaptation dynamics. Performance evolution during Test-Time Adaptation (TTA). Healthy subjects align in ∼1 epoch, while mTMD participants require ∼40 epochs, quantifying the biological domain shift. These specific trajectories represent post-hoc evaluations across the full target cohorts. Predictive performance reported in Table 1 utilizes strictly isolated calibration subsets to determine epoch hyperparameters, preventing data leakage. **f-g.** Uniform Manifold Approximation and Projection (UMAP)^49^ visualization of latent embeddings before and after Test-Time Adaptation (TTA) for Control (**f**) and mTMD (**g**) cohorts. TTA aligns the previously scattered mTMD embeddings into a coherent pain manifold. Spearman’s correlation between the UMAP components shown in (**f**) and (**g**) and the subjective pain ratings are shown under component axes. High correlations in the adapted latent space confirm that the model’s internal representation explicitly encodes pain intensity gradients. Statistical significance for Spearman’s rank correlations is indicated as follows: * P < 0.05, ** P < 0.01, *** P < 0.001; n.s., not significant.

**Table 1:** Performance comparison of LENS against classical benchmark models.

| Test Set | LENS |  | PLS-Preregistered |  | Linear-Spectral Shift |  | Linear-EMG |  | XGBoost-EMG |  |
| --- | --- | --- | --- | --- | --- | --- | --- | --- | --- | --- |
|  | Left | Right | Left | Right | Left | Right | Left | Right | Left | Right |
| *Training on Control-train* |  |  |  |  |  |  |  |  |  |  |
| Control-test<br>(*p*-value) | **0.52**<br>(0.0001) | **0.48**<br>(0.0005) | 0.22<br>(0.07) | -0.08<br>(0.70) | 0.07<br>(0.33) | 0.16<br>(0.15) | 0.04<br>(0.38) | -0.10<br>(0.76) | 0.13<br>(0.19) | -0.10<br>(0.75) |
| mTMD<br>(*p*-value) | **0.61**<br>(0.0001) | **0.48**<br>(0.0005) | **0.28**<br>(0.02) | 0.15<br>(0.15) | -0.15<br>(0.86) | 0.06<br>(0.33) | -0.09<br>(0.74) | 0.00<br>(0.49) | 0.16<br>(0.12) | **0.31**<br>(0.009) |
| *Training on mTMD* |  |  |  |  |  |  |  |  |  |  |
| mTMD*<br>(*p*-value) | **0.46±0.2**<br>(0.0003) | **0.40±0.2**<br>(0.0028) | -0.08±0.3<br>(0.68) | -0.15±0.2<br>(0.84) | 0.06±0.3<br>(0.35) | -0.14±0.3<br>(0.82) | -0.16±0.2<br>(0.86) | -0.03±0.1<br>(0.59) | -0.05±0.3<br>(0.62) | -0.23±0.4<br>(0.94) |
| Control-train<br>(*p*-value) | **0.21**<br>(0.034) | **0.20**<br>(0.045) | **0.19**<br>(0.045) | **0.38**<br>(0.0005) | -0.09<br>(0.78) | 0.10<br>(0.19) | -0.18<br>(0.95) | **0.21**<br>(0.03) | 0.12<br>(0.15) | 0.08<br>(0.23) |
| Control-test<br>(*p*-value) | -0.01<br>(0.53) | -0.19<br>(0.89) | **0.32**<br>(0.01) | -0.09<br>(0.73) | -0.07<br>(0.69) | 0.08<br>(0.29) | **0.33**<br>(0.01) | 0.05<br>(0.37) | 0.18<br>(0.12) | 0.07<br>(0.32) |
| *Training on All Subjects* |  |  |  |  |  |  |  |  |  |  |
| All*<br>(*p*-value) | **0.36±0.1**<br>(0.0001) | **0.35±0.1**<br>(0.0001) | **0.17±0.1**<br>(0.01) | **0.12±0.1**<br>(0.048) | -0.03±0.2<br>(0.66) | -0.05±0.2<br>(0.76) | 0.07±0.2<br>(0.16) | **0.15±0.2**<br>(0.02) | **0.30±0.2**<br>(0.0002) | 0.09±0.2<br>(0.12) |

Spearman’s rank correlation ( $\rho$ ) between predicted and true pain ratings within and across domains for the left and right masseters. LENS consistently outperforms traditional feature-engineering baselines (PLS, Linear-Spectral Shift, Linear-EMG, and XGBoost-EMG), demonstrating that deep sequence modelling is required to decode continuous MSK pain. The performance gap is particularly evident in zero-shot cross-domain transfer (Control-train → mTMD), where classical models fail to generalize. For 5-fold Cross-Validation (CV) experiments (indicated by an \*), results are reported as Mean ± SD. For cross-domain transfer, results are reported as single correlation coefficients. Permutation test *p*-values (10,000 iterations) are shown in parentheses. Significant correlations ( $p < 0.05$ ) are bolded.

To determine if this predictive capability extends to chronic MSK pain disorders, we trained and tested the model on the mTMD cohort using a stratified 5-fold cross-validation scheme. Despite the potential variability introduced by chronic myalgia, such as altered motor recruitment strategies, splinting, or protective guarding^36,37,47^, the model effectively predicted post-MVC pain intensity, yielding rank correlations comparable to the healthy cohort (left: ρ=0.46 ± 0.22; right: ρ=0.40 ± 0.20; Table 1). A unified model trained through 5-fold cross-validation on all 183 participants, agnostic to pathological status, retained significant but modest predictive power (left: ρ=0.36 ± 0.10; right: ρ=0.35 ± 0.11; p=0.0001), the signal diluted by the distinct physiological states of the two groups.

We benchmarked LENS against four classical pipelines, i.e., standard machine-learning models built on handcrafted signal features rather than learned representations. These included a pre-registered partial least squares (PLS) model using 23 time-domain, spectral, wavelet, non-linear (sample entropy), and hemodynamic features (Table S2; see *Methods*). Across all primary evaluation settings, LENS substantially outperformed the classical baselines (Table 1). Notably, while hand-crafted features and traditional models like PLS demonstrated marginal predictive power in specific cases (e.g., trained on mTMD, tested on Control-train; ρ=0.19–0.38), in other evaluation settings they either collapsed or were substantially outperformed by LENS (Table 1). This indicates that handcrafted summary statistics oversimplify the physiological signal, and that deep temporal modelling was required here for extracting a robust, generalizable pain signature.

To evaluate the translational utility of LENS for binary clinical endpoints, we assessed its capacity to distinguish moderate-to-severe pain (VAS≥45/100)^48^ from mild or no pain. LENS demonstrated robust diagnostic discrimination, achieving an Area Under the Receiver Operating Characteristic (AUROC) curve of 0.83 (left) and 0.87 (right) in the Control-test cohort and 0.86 (left) and 0.82 (right) in the unseen mTMD cohort (Fig. 2c). Calibration analysis revealed that while the model slightly over-predicts pain in healthy controls and under-predicts pain in mTMD cohort, predictions monotonically track observed pain gradients across the clinical spectrum (Fig. S1a). Furthermore, Precision-Recall analysis confirmed predictive stability above baseline prevalence in both cohorts (Fig. S1b and c). These operating characteristics are consistent with use for group-level stratification.

### A signature learned from healthy muscle transfers to chronic pain

We tested whether the model trained on healthy controls predicted pain in the unseen mTMD cohort, and it did so successfully (left: ρ=0.61; right: ρ=0.48, p < 0.001; Table 1; Fig. 2d). This performance was comparable to models trained directly on mTMD data. However, we observed an asymmetric transfer where healthy to mTMD transfer was highly effective, while the reverse, mTMD to healthy, yielded substantially lower performance (ρ=0.20–0.21; Table 1).

### Label-free adaptation aligns individual physiology

Test-time adaptation (TTA) was critical to this transfer. We measured predictive performance as a function of TTA epochs (Fig. 2e). While healthy subjects required only a single epoch of adaptation to reach optimal performance, the mTMD cohort required substantially more, peaking at approximately 40 epochs.

To understand the mechanism driving this disparity, we analysed the evolution of the model’s latent embeddings for both the healthy controls and the mTMD participants. Prior to adaptation, the latent representations of mTMD participants were unstructured, reflecting the profound biological distance between healthy and pathological states. Following TTA, the embeddings aligned into coherent manifolds. When evaluating the principal components of these adapted spaces, we found that low and high pain states were clearly separable and strongly correlated with subjective pain intensity in both the control (Fig. 2f) and mTMD cohorts (Fig. 2g). Crucially, this effect was not driven by hardware differences; the improvement with increased TTA epochs was observed regardless of whether recordings from mTMD participants were obtained with Device A or Device B (Fig. S2). This indicates that TTA reduces the individual-specific variance that obscures a conserved representation, rather than merely calibrating for sensor characteristics (see also Fig. S3 for evolution of encoder representations through training stages).

NIRS recordings were available for every participant, so the adaptation reported above minimized the full pre-training objective, which contains both an sEMG and a NIRS reconstruction term. Deployment, however, assumes sEMG alone. We therefore repeated the adaptation using the sEMG term only, holding all other settings fixed. On the primary left masseter, performance was unchanged within the healthy domain (Control-test ρ=0.51, compared with 0.52) and on transfer to mTMD (ρ=0.62, compared with 0.61). The secondary right masseter was slightly lower in the transfer setting only (ρ=0.43, compared with 0.48; Table S1). Adaptation therefore does not depend on the hemodynamic signal, and NIRS is required only to pre-train the encoder.

### High-frequency motor activity carries the pain signature

To identify the features driving LENS predictions, we performed attribution analysis. Attention weights within the conformer architecture revealed divergent temporal strategies between groups: the model trained on healthy controls attended predominantly to inter-trial rest periods, whereas the models trained on mTMD attended predominantly to the MVC phase itself (Fig. 3a).

**Fig. 3:**
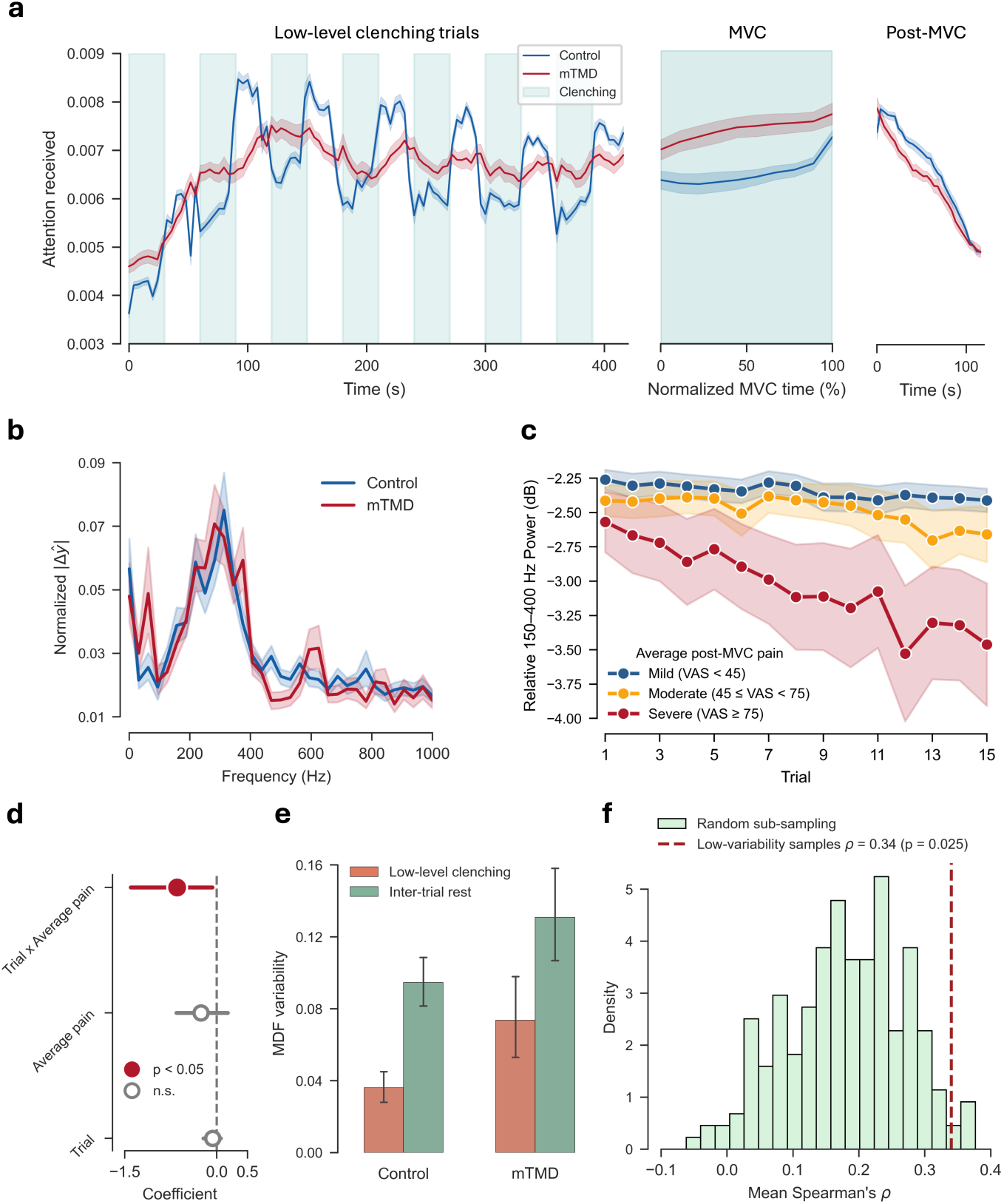
Physiological interpretability of LENS. **a.** Temporal attention weights. The Conformer attention map (averaged over heads) reveals distinct strategies: models trained on healthy controls prioritize inter-trial rest periods (recovery), whereas models trained on mTMD prioritize the MVC phase (exertion). **b.** Spectral occlusion analysis. Relative importance of EMG frequency bands. The model relies heavily on the 150–400 Hz range for pain prediction. **c.** Observed temporal evolution of relative 150–400 Hz EMG band power across 15 clenching trials, stratified by post-MVC subjective pain severity: mild (VAS<45), moderate (45≤VAS<75), and severe (VAS≥75). Shaded regions represent the standard error of the mean (SEM). Higher pain severity corresponds to a steeper spectral decline. **d.** Forest plot of LMM fixed-effect coefficients (trial, pain, and trial × pain interaction) demonstrating that subjective pain significantly accelerates the decline of high-frequency power. Points and error bars represent the mean and 95% confidence intervals derived from 5,000 subject-level block bootstrap iterations. **e.** Motor variability across cohorts. Within-subject variability of the median frequency (MDF) across the final seven trials is significantly elevated in the mTMD cohort compared to healthy controls during both inter-trial rest (p=0.005, two-sided Mann-Whitney U test) and low-level clenching (p=0.001). Error bars denote the 95% confidence intervals. **f.** Random sub-sampling control for zero-shot transfer. The histogram represents the empirical null distribution of predictive performance (mean bilateral Spearman’s ρ on healthy controls) generated from 200 models trained on randomly selected, size-matched mTMD subsets (n=31). The dashed vertical line marks the performance of the model trained strictly on the low-variability mTMD subset (n=31, MDF variability ≤ 0.2). Pruning high-variability pathological signatures significantly improves generalization compared to random sampling (empirical p=0.025).

Spectral occlusion analysis revealed that the model’s predictions relied heavily on the 150–400 Hz frequency band (Fig. 3b), despite the bulk of raw EMG power residing below 100 Hz (Fig. S4). To empirically ground this machine-learned feature in muscle physiology, we extracted the relative signal power in the 150–400 Hz band and tracked its temporal evolution across the 15 clenching trials. Stratifying these trajectories by pain severity revealed that higher subjective pain was associated with a steeper decline in high-frequency power over time (Fig. 3c). This is reflected in a significant pain × trial interaction in a linear mixed-effects model with participant-level random intercepts and slopes (p=0.025; Fig. 3d). Crucially, a parallel model evaluating clinical diagnosis (mTMD vs. healthy control)—rather than subjective pain—found no significant group × trial interaction (p=0.252; Table S3) indicating that the temporal decline in high-frequency power tracks pain intensity specifically, rather than disease status. mTMD participants did show lower high-frequency power overall (p=0.028; Table S3), but this static difference did not extend to the trial-wise dynamics captured by the pain-tracking effect.

### Motor variability explains the asymmetry of transfer

To test a potential source asymmetric transferability, we hypothesized that idiosyncratic motor adaptations in chronic MSK pain act as physiological noise, obscuring the canonical pain signature during model training. We quantified within-subject motor variability using the coefficient of variation of the median frequency (MDF) across the final seven trials (trials used as input for LENS; see Methods). Confirming our hypothesis, participants with mTMD exhibited significantly higher motor variability than healthy controls during both inter-trial rest (Mann-Whitney U=4397.0, p=0.005) and low-level clenching (U=4524.0, p=0.001; Fig. 3e).

To test whether these highly variable pathological signatures were the source of the transfer failure, we applied a data-pruning strategy: excluding 23 mTMD participants with excessive motor variability (MDF variability > 0.2 on either side during rest or task^50^) and retraining the model on the remaining subset (n=31). This model achieved a substantial improvement in generalization to the healthy Control-train cohort (ρ=0.34 bilaterally). Against an empirical null of 200 models trained on size-matched random mTMD subsets (n=31; *Methods*), the pruned model outperformed 97.5% of draws (empirical p=0.025; Fig. 3f), excluding sample-size reduction as the explanation. This indicates that the canonical pain signature is conserved in mTMD but is heavily masked by pathology-specific motor variance.

### Predictions track pain rather than fatigue

Because the task is fatiguing by design, subjective fatigue is the principal confound for a pain-specific claim. In the healthy Control-test cohort, fatigue explained under 6% of the rank variance in reported pain, and LENS predictions were themselves unrelated to fatigue (ρ=0.10–0.18, p > 0.05; Fig. 4a). The association between LENS predictions and pain was undiminished after partialling out fatigue (left: partial ρ = 0.50, p=0.0003; right: 0.48, p=0.0007), while the reciprocal association between LENS predictions and fatigue, after partialling out pain, was absent (ρ=0.07 and 0.02; both p > 0.6). This double dissociation indicates that LENS tracks pain specifically, not fatigue. In the mTMD cohort, however, pain and fatigue ratings were near-collinear (ρ=0.79–0.81), so LENS predictions correlated with both fatigue (ρ=0.54 and 0.51) and pain. Because controlling for one removed most of the variance associated with the other, neither partial correlation is interpretable, and we report them for completeness only (Table S4).

**Fig. 4:**
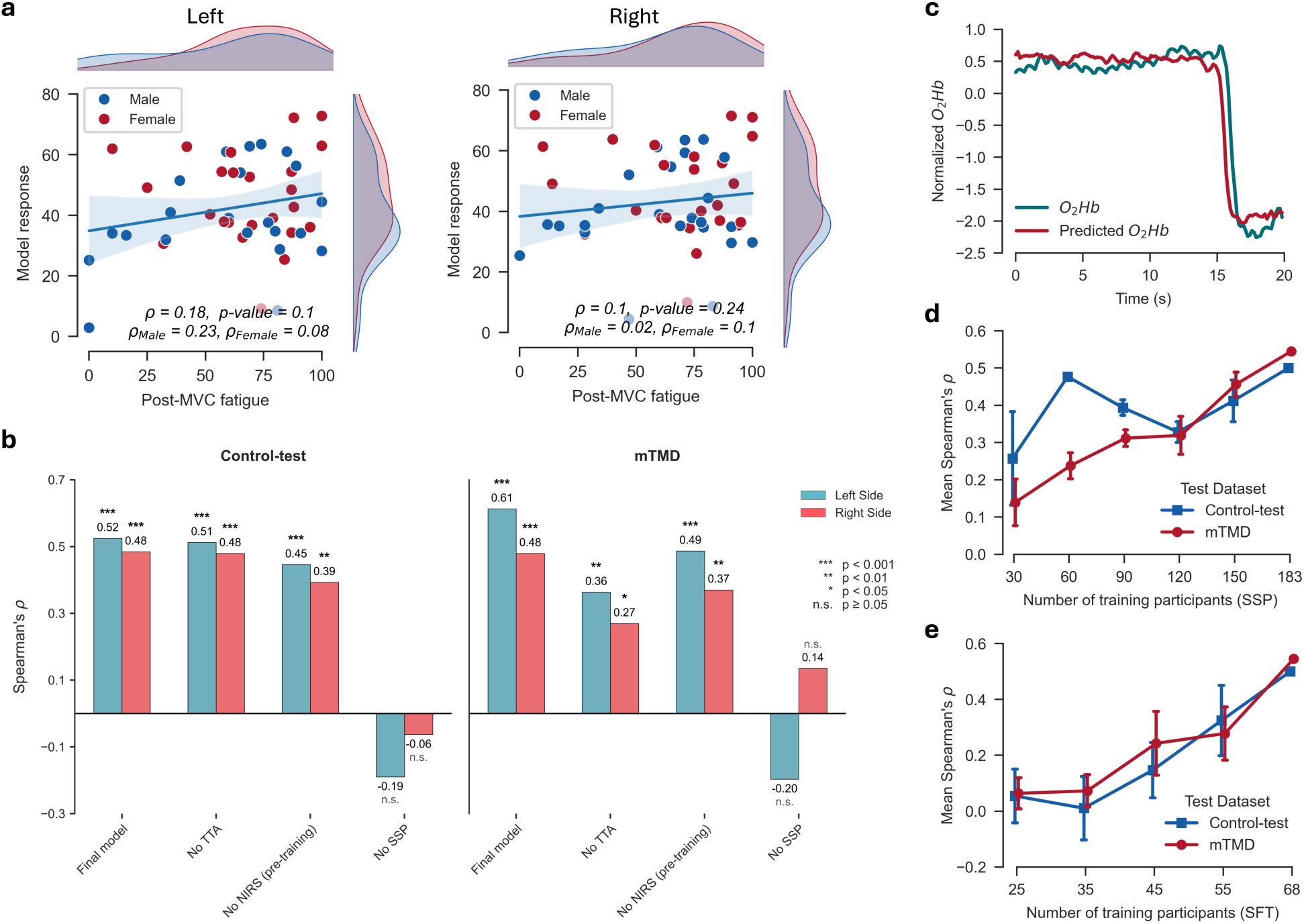
Model specificity, component necessity, and data scaling. **a.** Specificity to pain. LENS predicts subjective pain but shows no significant correlation with fatigue ratings (ρ<0.2) in the Control-test cohort, disentangling nociception from exhaustion. **b.** Ablation study. Spearman’s correlation drops significantly without Self-Supervised Pre-training (SSP) or the NIRS reconstruction objective (No NIRS) during pre-training, supporting the contribution of the hierarchical, cross-modal approach. **c.** Cross-modal reconstruction. A representative example of the EMG encoder reconstructing the O2Hb (oxygenated hemoglobin) signal during self-supervised pre-training, demonstrating learned neurovascular coupling. **d-e.** Data scaling laws for SSP (**d**) and Supervised Fine-Tuning (**e**). Pre-training achieves reasonable performance at ∼60 subjects, while fine-tuning benefits continuously from larger cohorts. Error bars show standard error across 5 randomized sub-samplings.

Component ablation supported our hierarchical training strategy (Fig. 4b). Removing the self-supervised pre-training (SSP) stage entirely collapsed predictive performance (ρ≈0). A more targeted ablation was performed by removing only the cross-modal reconstruction objective that aligns the sEMG encoder with concurrent NIRS-derived hemodynamics (Fig. 4c). This also resulted in degraded performance relative to the full model, although to a lesser degree than SSP ablation. This indicates that learning neurovascular coupling dynamics during pre-training benefits the encoder’s ability to interpret the electrical signal in a pain context, even though NIRS is unavailable at inference—supporting the translational relevance of this pre-training strategy for a single-channel sEMG deployment.

Finally, we analysed the scalability of our approach by evaluating model performance as a function of training dataset size. For the Self-Supervised Pre-training (SSP) stage, reasonable performance can be achieved at approximately 60 participants (Fig. 4d), whereas the Supervised Fine-Tuning (SFT) stage showed continuous improvement with more data and degraded rapidly when the sample size was halved (Fig. 4e). Multimodal acquisition is therefore required only for the ∼60-participant pre-training cohort; fine-tuning and deployment proceed with single-channel sEMG alone.

## Discussion

LENS identifies a peripheral signature of pain intensity that is preserved in chronic MSK pain despite disease-associated changes in motor physiology. A representation learned entirely from healthy muscle generalized to individuals with chronic mTMD (ρ=0.61), performing comparably to models trained directly on mTMD data, whereas transfer in the reverse direction was substantially weaker (ρ=0.21). This directional asymmetry is unlikely to reflect a fundamentally different peripheral encoding of pain in mTMD, as that would be expected to impair transfer in both directions. It was also not explained by training-set size or differences in pain-label distributions: mTMD-trained models predicted pain within the mTMD cohort itself (ρ=0.46), and the distributional shift was bidirectional (Fig. 1c). Instead, the asymmetry was driven by greater motor variability in mTMD. Trial-to-trial variability in EMG median frequency was elevated during both rest and clenching, and restricting the mTMD training set to the 31 least variable individuals improved transfer to healthy controls (ρ=0.34) beyond size-matched random subsets (empirical p=0.025). Together, these findings suggest that chronic mTMD preserves a shared peripheral pain signature while superimposing idiosyncratic motor adaptations that obscure it^36,37,47^. More broadly, this asymmetry suggests a strategy for developing generalizable peripheral pain biomarkers: discover representations in physiologically homogeneous healthy cohorts, then adapt and validate them in clinical populations where disease-related heterogeneity may otherwise obscure conserved pain-related physiology.

This observation has biological and methodological implications. Biologically, peripheral muscle activity in mTMD remains a correlate of reported pain intensity despite the central mechanisms that characterize chronic MSK pain^34,35^. Therefore, the peripheral pain signal is not altered in chronic pain but obscured by variability, potentially caused by pathological motor adaptations^36^, resulting in limited interpretability. Methodologically, this challenges the common assumption in biomarker discovery that clinically relevant models should be trained directly in patient populations. Our findings indicate that attempting to learn both pain and pathology simultaneously can direct the model to preferentially learn pathological motor patterns at the expense of the more generalizable pain signal.

We deviated from conventional sEMG analysis and took advantage of the flexibility of deep neural networks to recover this obscured pain signal. Classical pipelines built on summary features yielded weak and inconsistent correlations (Table 1). LENS instead learns from the full time– frequency representation and models how it evolves across the task, so the predictions are informed by the cumulative history of exertion. Furthermore, the encoder is pre-trained to reconstruct concurrent NIRS hemodynamics, a relationship tightly coupled in healthy muscle^42^ but disrupted in myalgia^12,24,43,44^. Ablating either component degraded performance, and removing self-supervised pre-training collapsed it (Fig. 4b). This comparison implies that physiological biomarkers may not be found in single measurements, but rather in how a signal evolves and relates to physiological state over time.

LENS’s spectral profile points to the importance of high-frequency EMG patterns in pain prediction. Although masseter EMG power resides predominantly below 100 Hz (Fig. S4), occlusion analysis revealed that LENS relies disproportionately on the 150–400 Hz band (Fig. 3b), to which fast-twitch (type II) motor units contribute most^51,52^. Relative 150–400 Hz power declined more steeply across the clenching trials as reported pain increased (pain × trial interaction, p=0.025), while substituting clinical diagnosis for pain intensity produced no group × trial interaction (p=0.252). Therefore, the accelerated decline is associated with pain experienced after maximal exertion, not diagnosis. This may reflect early exhaustion or inhibition of highly fatigable type II units through localized metabolic stress or nociceptive reflex inhibition. However, our data do not distinguish these. The attention profile of LENS shows that models trained on mTMD shifted attention toward maximal exertion, whereas healthy-trained models weighted inter-trial rest, the phase in which recovery capacity is expressed^22,23^. Therefore, the physiological context carrying pain information shifted with disease state, suggesting that a biomarker may need to be sought in different temporal windows depending on disease state, rather than at one fixed time.

From a translational perspective, acquisition burden and inter-individual variability are the most important barriers limiting clinical implementation. LENS minimizes acquisition burden by internalizing the hemodynamic information during pre-training. NIRS, the more costly and technically demanding modality, is needed only to pre-train the encoder, achievable with approximately 60 participants (Fig. 4c). Deployment afterward requires a single sEMG channel per muscle. Inter-individual variability is addressed by test-time adaptation. This adaptation is unsupervised, uses roughly 5 min of the individual’s own unlabelled signal, and completes in under 13 s on a single CPU core. The number of adaptation steps each group needs to reach peak performance indicates how far the testing cohort’s physiology has departed from the training cohort’s manifold. LENS could complement self-report by tracking physiological response to treatment or by supporting patient phenotyping. It could also serve as a pharmacodynamic endpoint in early analgesic development, where efficacy is currently inferred from subjective ratings alone.

Our study has limitations. The association between predicted and reported pain is moderate (ρ ≈ 0.5), which is likely a ceiling imposed by the ground truth. Pain is a biopsychosocial experience shaped by mood, memory and social context^53–56^, none of which is directly reflected in a muscle signal. LENS is therefore a physiological correlate that complements self-report rather than an alternative to it. The pain decoded here was experimentally evoked in every cohort, including chronic mTMD, so we do not establish that LENS tracks spontaneous clinical pain^33^ or that it would work during unconstrained daily activity. The protocol is also active and requires a supervised clenching sequence, which is not equivalent to background monitoring. Generalization was demonstrated across cohorts, individuals, pathological status and two device families, but within a single muscle, at one centre, and in a demographically narrow sample (mean age mTMD: 26.15±5.25 years, Control-test: 23.9±6.21 years; 109 of 183 female across cohorts) with predominantly mild-to-moderate mTMD. Finally, to the best of our knowledge, no publicly available dataset pairs experimentally evoked muscle pain with sEMG recordings from the corresponding muscle and self-report pain ratings, which precluded external validation of the model^57–63^. External validation will therefore require prospective data collection. We outline a protocol for this in Supplementary Note 2 and release frozen model weights to enable independent replication.

Our results indicate that peripheral muscle carries a representation of MSK pain intensity that survives chronic pathology and can be recovered from a single sEMG channel per muscle, once pathology-specific motor variance is separated from it. LENS learned this representation more readily from healthy than from patient muscle. Therefore, discovery cohorts for peripheral pain biomarkers may be better assembled from healthy volunteers, with patient data reserved for evaluation and adaptation. How broadly this principle generalizes remains open, and establishing clinical usefulness will require external validation, testing in muscles other than the masseter, and evaluation against spontaneous pain.

## Methods

### Participants

Participants were recruited from the local community through flyers and advertisements on social networks linked to an online survey (REDCap) hosted on the local university server. The survey included demographic information and the DC/TMD Pain Screener^64^, a validated questionnaire that evaluates TMD symptoms over the past 30 days (e.g., presence of pain in the jaw or temple area exacerbated by jaw function). The screener has high specificity and sensitivity for detecting TMD when the cumulative score is ≥ 3. Participants with scores ≥ 3 were invited to undergo a clinical examination performed by a single examiner (IC), who has clinical expertise in TMD. The examination included a detailed clinical history, assessment of mandibular range of motion, evaluation of dental status, and palpation of the masticatory muscles. The clinical assessment was performed according to the DC/TMD^65^ and the guidelines outlined by the International Network for Orofacial Pain and Related Disorders Methodology (INfORM), International Association for Dental, Oral, and Craniofacial Research (IADR) (https://inform-iadr.com/). Participants diagnosed with TMD myalgia, myofascial pain, or myofascial pain with referral were included in the study. The presence of articular disorders, such as disc displacement with reduction or TMJ arthralgia, was not considered an exclusion criterion. Healthy controls were recruited from individuals who scored 0 on the DC/TMD Pain Screener and did not report any TMD symptoms.

We collected data from 54 individuals with myogenous mTMD (40F; 26.15±5.25 years) and two independent cohorts of healthy controls for training (n=81; 45F; 25.96±5.9 years) and test (n=48; 24F; 23.9±6.21 years). All participants consented to the procedures described, which were reviewed and approved by the University of Toronto Human Research Ethics Board (protocol #39063). Participants were financially compensated.

Exclusion criteria for control participants were: (1) presence of TMD and/or orofacial pain; (2) in treatment with medications affecting the activity of masticatory muscles or the sensory system; (3) extensive dental prostheses (>6 teeth); (4) undergoing dental/orthodontic treatment; (5) neurological disorders (e.g., epilepsy, seizures, Alzheimer’s disease, Parkinson’s disease, multiple sclerosis, etc.) and/or using medications for the management of such neurological disorders, autoimmune, endocrine, cardiovascular disease, cancer, diabetes. Exclusion criteria for mTMD participants were: (1) extensive dental prostheses (>6 teeth); (2) undergoing dental/orthodontic treatment; (3) neurological disorders (e.g., epilepsy, seizures, Alzheimer’s disease, Parkinson’s disease, multiple sclerosis, etc.) and using medications for the management of such neurological disorders; (4) non-compensated endocrine conditions; (5) autoimmune conditions, cardiovascular disease, cancer, diabetes; (6) drug or alcohol abuse disorders; (7) pregnancy.

mTMD participants completed the Graded Chronic Pain Scale (GCPS) as part of the DC/TMD assessment. The GCPS evaluates pain intensity and pain-related disability over the past 30 days, categorizing individuals into hierarchical grades of pain severity: grade 0 (pain-free), grade 1 (low intensity pain, none-low pain-related disability), grade 2 (high intensity pain, none-low pain-related disability), grade 3 (moderately limiting), and grade 4 (severely limiting). The distribution of GCPS grades for the mTMD cohort is shown in Supplementary Fig. S5. mTMD participants reported 96±65 days of facial pain in the last six months, facial pain of 3.0±2.2 (0–10 scale) on the day of the experiment before starting the tasks, average facial pain of 3.9±1.8 (0–10 scale) in the last 30 days, and worst facial pain of 6.3±2.2 (0–10 scale) in the last 30 days.

### EMG biofeedback

Before the experimental task (see below), participants’ maximum voluntary contraction (MVC) was recorded. Participants sat on a chair with their head unsupported. The skin on the masseter on each side was cleaned by rubbing isopropyl alcohol. One concentric electrode (30 mm, OT Bioelettronica) (or bipolar 24 mm electrodes, OT Bioelettronica) was placed on the masseter muscles, on the most prominent part of the muscle belly identified during maximum voluntary contraction, along the muscle long axis. The electrode was connected to an EMG device connected to a computer via Bluetooth. We have used two different EMG devices during data collection: Due (“EMG device A”; 2048 Hz sampling frequency) and Due+ (“EMG device B”; 2000 Hz sampling frequency) (OT Bioelettronica, Turin, Italy). A NIRS device (Portalite Mini, Artinis, Elst, the Netherlands) was positioned horizontally on the upper part of the masseter muscle beneath the zygomatic arch. The EMG electrode was placed beneath the NIRS optodes, on the lower part of the masseter muscle. The NIRS device has three optodes, which sample at 16, 21, and 26 mm depths. Oxygen tissue saturation (StO_2_), deoxygenated hemoglobin (HHb), and oxyhemoglobin (O_2_Hb) signals from NIRS were used in this study.

Real-time visual feedback of EMG amplitude was provided from the right masseter only; participants were not informed which side the display was derived from. Prior to the task, each participant’s MVC was measured by asking them to bite as hard as possible in maximum intercuspation for five seconds, repeated three times. The MVC reference used to define the 20– 30% target range was likewise computed from the right-side signal, so the contraction level of the left masseter was neither displayed to the participant nor normalized to a left-side maximum. The MVC amplitude is calculated by taking the average of the samples the computer receives over each time window of 0.0625 seconds and then taking the maximum of these averages over five seconds. This method of smoothing helps to obtain a more reliable measure by attenuating the effect of very transient, biologically irrelevant EMG oscillations. Median of the three recorded values was then used as the MVC. Participants were asked to verify the registration of their MVC by clenching their teeth at different force levels to ensure that the registration matched their perception. EMG visual feedback was then constructed using the software so that participants could engage in alternated tooth clenching and rest activities at specific times and target specific ranges of MVC and duration (see experimental task), while looking at a computer screen.

### Experimental task

An experimental task was designed to record the masseter functional response to a repeated and low-level tooth clenching. NIRS and EMG signals from right and left masseter were collected throughout the task (Figure 1b). Visual analogue scales for pain and fatigue (VAS^45^, 100 mm; left anchor: “no pain/fatigue”; right anchor: “the worst pain/fatigue I can imagine”) were used to collect pain reports at the masseter and temporalis bilaterally, at the end of each tooth clenching trial.

The experiment started with 135 seconds of rest. Thereafter, participants engaged in a repetitive tooth clenching task in which they had to target 20% to 30% MVC for 30 seconds, followed by 30 seconds of rest (see EMG biofeedback). This cycle of 30 sec clenching and 30 sec rest were repeated 15 times. Thereafter, participants rested for 195 seconds and then clenched their teeth at their MVC until failure. Pain and fatigue ratings were also collected after this last clenching exercise (post-MVC pain), which was followed by rest (∼2 min).

### Data pre-processing

Synchronized sEMG and fNIRS recordings were aligned to a common start time and trimmed to match durations. sEMG Processing: To standardize acquisitions across devices, sEMG signals were resampled to 2000 Hz and band-pass filtered (40–850 Hz) using a zero-phase 4th-order Butterworth filter. Time–frequency features were extracted via log-spectrograms computed with a 32 ms analysis window and a 20 ms stride. fNIRS Processing: We retained individual optode channels for oxygenated (O_2_Hb) and deoxygenated (HHb) hemoglobin without spatial averaging. Signals were baseline-normalized relative to the initial two minutes of recording, and their first temporal derivatives were computed and appended to the feature set. Feature alignment: All feature sequences and task labels were temporally aligned to the 50 Hz resolution defined by the spectrogram stride.

### Model Architecture and Training Strategy Deep Learning Architecture

We developed a hierarchical sequence modelling framework to map high-dimensional sEMG log-spectrograms to subjective pain ratings. The model, comprising approximately 2.2 million trainable parameters, processes inputs consisting of concatenated log-spectrograms from left and right masseters (33 frequency bins × 2 channels). The architecture proceeds in three stages. First, a local feature encoder, consisting of a 5-layer Time-Depth Separable (TDS) convolutional network^46^, extracts local temporal features while performing an 8x temporal downsampling. Second, a convolutional downsampler comprising two 1D convolutional layers projects the encoder output into the transformer latent dimension, performing a further 25x temporal reduction. This sequence is concatenated with a 3D one-hot embedding representing the task phase (baseline, trial, or MVC) and processed by a Global Sequence Modeler—a 4-layer Conformer (convolution-augmented transformer)^39^. The Conformer utilizes 4 attention heads (configured as 3 local and 1 global) and models relative position information using a simplified T5-style bias (16 buckets, maximum distance 256, shared across heads)^66^. Finally, a learned class token (CLS) is prepended to the sequence; its output embedding drives a Multi-Layer Perceptron (MLP) regression head to predict post-MVC pain intensity.

### Training Strategy

To address the scarcity of labelled clinical data and the heavy-tailed distribution of pain ratings, we employed a three-stage training strategy. Data augmentation was applied throughout all stages, utilizing SpecAugment^67^ for sEMG (time and frequency masking) and additive jitter noise for NIRS.

Stage 1: Cross-Modal Self-Supervised Pre-training (SSP). We pre-trained the encoder using a masked latent-space modelling objective on random 20-second signal chunks to capture local dynamics using SmoothL1 loss^68^ and cosine similarity with a batch size of 24 (8 per GPU). Let *X* ∈ ℝ*^F^*^×*T*^ represent the input log-spectrogram and *Y* ∈ ℝ*^C^*^×*T*^ represent the concurrent NIRS hemodynamic signals. We generated binary masks *M_EMG_* and *M_NIRS_* to select time-frequency regions for reconstruction. To account for the disparate temporal dynamics of the two modalities, we applied differential masking strategies. For the stochastic, rapidly varying sEMG signal, we used a mask probability of p=0.5 with a window size of ∼200 ms. Conversely, given the slow physiological time constants of the hemodynamic response, we employed a significantly larger window for NIRS (∼3 s, p=0.45); this forces the network to infer long-range metabolic trends rather than interpolating local noise.

The model was optimized to reconstruct the values of the masked regions only. The input to the model was the EMG masked view *X̃* = *X*⊙*M_EMG_*, the target for cross-modal reconstruction was NIRS masked view *Ỹ* = *Y*⊙*M_NIRS_* and the joint loss ℒ*_SSP_* was calculated as:

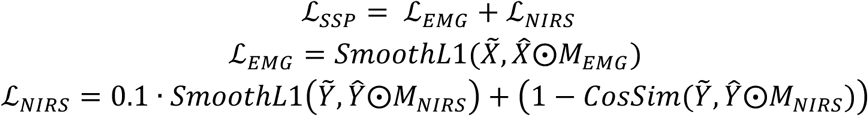

where *X̂* and *Ŷ* are the model outputs, and CosSim(⋅) is the cosine similarity along the temporal dimension. This cross-modal objective ensures the encoder captures directional hemodynamic trends coupled with neuromotor activity.

Stage 2: Supervised Fine-Tuning (SFT). The pre-trained encoder and randomly initialized Conformer were fine-tuned end-to-end on the labelled datasets. For this phase, inputs were constructed by concatenating the last 7 clenching trials (trials 9–15), the MVC phase, and the two-minute post-MVC window. This restriction to the final seven trials was empirically motivated by the pain reporting distribution; by trial 9, approximately 50% of the training cohort reported a pain intensity exceeding a minimum distinct perception threshold (5/100 VAS)^48^, thereby maximizing the signal-to-noise ratio of the labels (Fig. S6). We employed early stopping with a patience mechanism based on the validation loss (calculated on a 15% split of the training data). Labels *_y_* were log-transformed and z-scored to stabilize gradients:

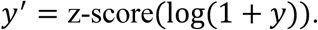

Standard Regression: For cohort-specific models (control/mTMD), we minimized the SmoothL1 loss:

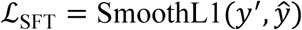

Probabilistic Regression: For the unified model trained on all subjects, we addressed the non-normal, bimodal label distribution by replacing the MLP head with a Mixture of Two Beta Distributions. The network outputs parameters *φ* ∈ ℝ^5^ per side, mapping to a mixing weight *w* = *σ*(*φ*_0_) and shape parameters {*α_k_*, *β_k_*} = softplus(*φ*_1..4_) + *ε* for components *k* ∈ {1,2}. We minimized the Negative Log-Likelihood (NLL):

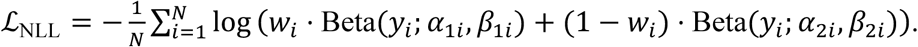

Point predictions were obtained via the weighted mean of the expected values:

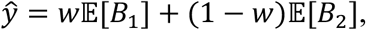

where

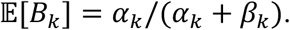

Stage 3: Test-Time Adaptation (TTA). To mitigate domain shifts caused by pathological differences or electrode placement variations, we applied test-time adaptation for each test subject. For a test subject with unlabelled input *X*_test_, we updated the encoder parameters *θ*_enc_ to minimize the self-supervised objective from Stage 1:

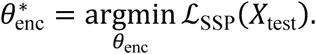

This adaptation aligns the feature space of the test subject with the training manifold prior to inference, while the Conformer and downstream regression head remain frozen.

Because NIRS was recorded in all participants, the adaptation results reported in the main text minimize the full Stage 1 objective, ℒ*_SSP_* = ℒ*_EMG_* + ℒ*_NIRS_*. Deployment requires adaptation from sEMG alone, so we repeated the procedure minimizing ℒ*_EMG_* only, with all other settings unchanged. Results are given in Supplementary Table S1.

During TTA, the model was updated using 16 randomly selected 20-second signal chunks from each subject with a batch size of 4. To strictly prevent data leakage and ensure rigorous evaluation, the optimal number of adaptation epochs was treated as a domain-level hyperparameter and determined prior to final inference using independent validation sets. We evaluated a discrete search space of {1,10,20,40} epochs, utilizing Spearman’s rank correlation (ρ) between predictions and ground-truth labels as the selection metric. For in-domain evaluations (e.g., training on Control-train and testing on Control-test), the optimal number of epochs was determined using the combined training and validation sets originally employed during the Supervised Fine-Tuning (SFT) stage. For cross-domain transfer evaluations (e.g., training on healthy controls and testing on mTMD), we randomly sampled a target-domain calibration set consisting of 6 subjects (approximately 10% of the mTMD cohort). This calibration set was used exclusively to determine the optimal adaptation trajectory for the target domain and was strictly excluded from the final held-out test set to maintain a true zero-shot inference paradigm with respect to labels.

AdamW^69^ optimizer (0.01 weight decay) with cosine learning-rate scheduling^70^ were used for training with an initial learning rate of 5 × 10^−5^ for SSP and SFT, and 1 × 10^−5^ for TTA. All deep learning architectures and training pipelines were implemented using PyTorch^71^. SSP and SFT (stages 1 and 2) were executed across three NVIDIA RTX A6000 GPUs utilizing PyTorch’s Distributed Data Parallel (DDP) while TTA was performed on a single GPU. To evaluate the practical feasibility of deploying the model in resource-constrained clinical settings without dedicated GPU infrastructure, TTA was benchmarked on a standard local workstation using a single CPU core (Apple Silicon M2 Ultra). TTA was completed in 325 ms for a single epoch (healthy participants) or 12.6 s for 40 epochs (participants with mTMD) on one CPU core.

### Model Interpretability and Visualization

Frequency Occlusion Analysis: To quantify the contribution of specific spectral bands, we employed a perturbation analysis. For each subject, a baseline feature vector *_X_*-was computed as the temporal mean of the first 120 seconds of the raw EMG features (baseline period). For each frequency bin *f*, we generated an occluded input sequence *X*^(*f*)^ by replacing the features at bin *f* (across both channels and all valid time steps) with the subject-specific baseline *_X_-_f_*. The importance of frequency *_f_* was quantified as the absolute deviation in the model’s prediction for the left side:

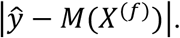

Attention Map Extraction: We extracted the temporal attention weights from the final Conformer layer to visualize the model’s focus. For a sequence of length *T*, we retained the attention matrix *A* ∈ ℝ*^T^*^×*T*^ (averaged over attention heads). We derived a 1D temporal importance curve *I* ∈ ℝ*^T^*^−1^ by computing the mean incoming attention from all non-CLS queries to a target token *k*:

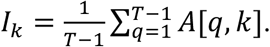

For analysis, these importance curves were phase-aligned using the downsampled phase IDs: the 7 trial windows were concatenated, the variable-length MVC segment was resampled to 10 points, and the post-MVC window was retained at native resolution (first 120 s).

Manifold Visualization: Latent space representations were visualized using Uniform Manifold Approximation and Projection (UMAP)^49^ with the following parameters with two components, n_neighbors of 15, min_dist of 0.1, and Euclidean distance metric.

### Classification Metrics and Physiological Signal Analysis

Classification Metrics and Calibration: To evaluate clinical operating characteristics, continuous VAS predictions were binarized using a VAS≥45/100 threshold, representing moderate-to-severe pain. Model discrimination was evaluated using the Area Under the Receiver Operating Characteristic curve (AUROC) and Precision-Recall (PR) curves for both Control-test and mTMD cohorts. Calibration was assessed by segmenting predictions into three terciles and plotting the mean predicted versus mean observed pain for each bin to evaluate monotonic tracking and conditional bias.

Physiological Signal Analysis (150–400 Hz Trajectories): To isolate the physiological drivers of the model’s spectral attention, raw EMG signals were resampled to 2000 Hz, band-pass filtered (40–850 Hz), and notch filtered to attenuate line noise and its harmonics (60, 120, 180, 240, 300, and 360 Hz). Power spectral density (PSD) was computed using Welch’s method. For each trial, relative high-frequency power was calculated as:

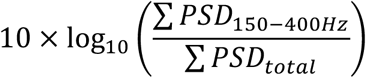

To statistically evaluate temporal dynamics, we utilized Linear Mixed-Effects Models (LMMs) via the statsmodels library^72^, incorporating random intercepts and random slopes for the trial variable grouped by subject. Predictors included scaled trial sequence (0–1) and scaled subjective pain (0–1) or group status. To derive rigorous 95% confidence intervals and mean coefficient estimates, we performed subject-level block bootstrapping (N=5000 iterations) to preserve intra-subject correlation structures.

Motor Variability and Random Sub-sampling Analysis: To quantify within-subject motor consistency, we calculated the Median Frequency (MDF) from the Power Spectral Density (PSD) of the EMG signals for the final seven clenching trials and their corresponding inter-trial rest periods. Variability was defined as the coefficient of variation (standard deviation divided by the mean) of the MDF across these seven windows. Group differences in MDF variability were evaluated using two-sided Mann-Whitney U tests.

To assess the impact of motor variability on cross-domain generalization, we established a pruning threshold, excluding 23 mTMD participants who exhibited an MDF variability > 0.2 on either the left or right side during either the clench or rest phases. The SFT and TTA steps were then performed from scratch on the remaining low-variability mTMD cohort (n=31) and evaluated on the healthy Control-train dataset. To control for the confounding effect of reduced sample size on model performance, we constructed an empirical null distribution via random sub-sampling. We trained 200 independent models using randomly selected mTMD cohorts of identical size (n=31), maintaining identical hyperparameters and early-stopping criteria. Model performance was summarized as the average Spearman’s ρ across the left and right masseters. An empirical p-value was calculated as the proportion of the 200 random iterations that yielded a mean ρ greater than or equal to that of the low-variability model.

### Classical Benchmark Models

To justify the deep learning architecture, we implemented four classical machine learning pipelines using handcrafted features. Steady-state trial windows were extracted to isolate motor and metabolic responses, removing 5-second transient buffers at contraction onset and offset. We extracted time-domain (RMS, Zero-Crossing Rate), frequency-domain (Mean/Median Frequency, High-Frequency Ratio, Dimitrov Fatigue Index), non-linear (Sample Entropy), wavelet (Discrete Wavelet Transform db5), and hemodynamic (StO_2_) features.

The baselines included: (1) a pre-registered Partial Least Squares Canonical regression model (5 features), (2) a Ridge regression Spectral-Shift model targeting classical median frequency literature (4 features), (3) a comprehensive Ridge regressor (23 features), and (4) a Gradient Boosted Decision Tree (XGBoost^73^) regressor (23 features). Independent models were trained for the left masseter (predicting self-reported left post-MVC pain VAS score) and right masseter (predicting right post-MVC pain VAS score) based on the features from the corresponding side. Hyperparameters were tuned via nested 10-fold cross-validation. Full mathematical formulations, the complete feature matrix, and hyperparameter grids are detailed in Supplementary Note 1. All the baseline models except XGBoost were implemented using the Scikit-learn library^74^.

### Statistical Analysis and Benchmarking

We evaluated performance across multiple benchmarks, including 5-fold cross-validation and cross-domain transfer (e.g., train on Control, test on mTMD). For cross-validation, folds were stratified by recording device (device A vs. B) to prevent hardware-driven confounding. Predictive performance was assessed using Spearman’s rank correlation (ρ), reported separately for the left and right sides. Statistical significance for cross-validation was determined via a stratified permutation test (shuffling labels within folds), while independent test set evaluations used a standard permutation test.

Partial Spearman correlations were computed, using the Pingouin package^75,76^, by rank-transforming each variable, regressing the covariate out of both the predictor and the outcome by ordinary least squares, and correlating the residuals; significance was assessed against a t distribution with n − 3 degrees of freedom. The proportion of rank variance in reported pain attributable to reported fatigue is given as the squared Spearman coefficient. Because partial correlation is uninformative when the covariate is collinear with the outcome, the primary specificity analysis was conducted in the Control-test cohort, in which pain and fatigue ratings are largely independent; the corresponding mTMD values are reported in Supplementary Table S4.

## Data availability

Anonymized preprocessed data from 152 participants who provided informed consent for data sharing are openly available at: https://borealisdata.ca/dataset.xhtml?persistentId=doi:10.5683/SP4/JCWT3H.

## Code availability

The full preprocessing, training, and evaluation pipeline codes for LENS and all the baseline models are openly available via GitHub at https://github.com/PedramMouseli/LENS.

## Author Contributions

P.M. and I.C. designed the experiment; P.M. performed the data curation, algorithm development, analysis, and visualization; I.C. performed the clinical screening of the mTMD participants; P.M., N.W.D., and S.S. collected the data; W.D.R., I.J., M.A., and N.A. provided insight and expertise for study methodology and reviewed the manuscript; P.M., M.M., and I.C. wrote the paper; and M.M. and I.C. supervised the research and provided guidance.

## Acknowledgements

This work was supported by a Canadian Institutes of Health Research Project Grant (N441030 -452593), Institute of Musculoskeletal Health and Arthritis Project Grant Priority announcement (RN401752), and the Bertha Rosenstadt Memorial Fund at Faculty of Dentistry, University of Toronto, as well as a Canada Fund for Innovation John R. Evans Leaders Fund – Funding for research infrastructure and the Ontario Research Fund – Research Infrastructure (IOF-37502). PM, SS, NDV was supported by the Harron Fund at the University of Toronto’s Faculty of Dentistry. SS was also supported by the American Association for Orthodontics, a University of Toronto Centre for the Study of Pain Scholarship and a Natural Sciences and Engineering Research Council Canada Graduate Scholarship—Master’s. MM holds a Canada Research Chair (Tier 2) in Pain NeuroImaging and is supported by a University of Toronto Centre for the Study of Pain Scientist Award.

