## Supplementary for "LENS: a transferable neuromuscular signature of musculoskeletal pain intensity"

[1]Centre for Multimodal Sensorimotor and Pain Research, Faculty of Dentistry, University of Toronto, Toronto, ON, Canada;
[2]Krembil Research Institute, University Health Network, Toronto, ON, Canada;
[3]University of Toronto Centre for the Study of Pain, Toronto, ON, Canada;
[4]Department of Physical Therapy, University of Toronto, Toronto, Canada;
[5]Interdepartmental Division of Critical Care Medicine, University of Toronto, Toronto, ON, Canada;
[6]KITE, Toronto Rehabilitation Institute, University Health Network, Toronto, ON, Canada;
[7]Osteoarthritis Research Program, Schroeder Arthritis Institute and Data Science Discovery Centre for Chronic Diseases, Krembil Research Institute, UHN, Toronto, ON, Canada;
[8]Departments of Medical Biophysics and Computer Science, and Faculty of Dentistry, University of Toronto, Toronto, ON, Canada;
[9]Institute of Neuroimmunology, Slovak Academy of Sciences, Bratislava, Slovakia;
[10]School of Digital Public Health, Mohamed bin Zayed University of Artificial Intelligence, Abu Dhabi, UAE;
[11]Department of Anesthesiology, Perioperative and Pain Medicine, Stanford School of Medicine, Stanford, CA, USA;
[12]Department of Pediatrics, Stanford University School of Medicine, Stanford, CA, USA
[13]Department of Biomedical Data Science, Stanford University, Stanford, CA, USA
[14]Immunology Program, Stanford University School of Medicine, Stanford, CA, USA
[15]Department of Dentistry, Mount Sinai Hospital, Toronto, ON, Canada.
[†]Contributed equally

**Table S1: Model performance with and without NIRS during test-time adaptation (TTA).**

| Test dataset | Full TTA | | EMG-only TTA | |
|---|---|---|---|---|
| | Left | Right | Left | Right |
| Control-test<br>*(p-value)* | **0.52**<br>*(0.0001)* | **0.48**<br>*(0.0005)* | **0.51**<br>*(0.0001)* | **0.48**<br>*(0.0004)* |
| mTMD<br>*(p-value)* | **0.61**<br>*(0.0001)* | **0.48**<br>*(0.0005)* | **0.62**<br>*(0.0001)* | **0.43**<br>*(0.001)* |

The model was trained on the Control-train dataset and tested on the Control-test and mTMD datasets. Adaptation used either both the EMG and NIRS reconstruction terms (Full TTA) or the EMG term alone (EMG-only TTA). All other settings, including the number of adaptation epochs, were held fixed. Left and Right refer to the left and right masseter, the left being the primary endpoint. Values are Spearman's rank correlation (ρ) between predicted and reported post-MVC pain, with permutation test p-values (10,000 iterations) in parentheses. EMG-only adaptation gives equivalent performance on the primary endpoint in both the healthy and the transfer setting. The secondary right masseter is unchanged in the healthy setting and lower on transfer to mTMD.

### Supplementary Note 1: Classical Benchmark Models and Feature Engineering

To isolate steady-state motor and metabolic responses while excluding movement transients at contraction onset/offset, for each 30s clenching and inter-trial rest period, a 5-second buffer was trimmed from both ends, yielding a 20-second steady-state window. The $StO_2$ signal was normalized by subtracting the mean of the baseline signal before the start of the task.

#### 1. Mathematical Formulation of Handcrafted Features

Features were extracted independently for the left ($[\mathrm{sEMG}_L, \mathrm{StO}_{2,L}]$) and right ($[\mathrm{sEMG}_R, \mathrm{StO}_{2,R}]$) masseters.

#### A. Time-Domain and Amplitude Features

- **Root Mean Square (RMS):** Quantifies gross motor unit firing amplitude during clenching and MVC trials:

$$\mathrm{RMS}^{(i)} = \sqrt{\frac{1}{N}\sum_{n=1}^{N} x[n]^2}$$

- **Zero-Crossing Rate (ZCR):** Evaluates the frequency of signal polarity changes per second, serving as a time-domain proxy for spectral shifts:

$$\mathrm{ZCR}^{(i)} = \frac{1}{T}\sum_{n=1}^{N-1} \mathbb{I}(x[n]\cdot x[n+1] < 0)$$

#### B. Spectral and Frequency-Domain Features

Power Spectral Density, $P(f)$, was estimated using Welch's periodogram method with non-overlapping Hamming windows of length $0.5f_s$.

- **Mean Frequency (MNF):** The centroid frequency of the PSD spectrum:

$$\mathrm{MNF}^{(i)} = \frac{\sum_k f_k\, P(f_k)}{\sum_k P(f_k)}$$

- **Median Frequency (MDF):** The frequency dividing the total spectral power into two equal halves:

$$\sum_{f_k=f_{\min}}^{\mathrm{MDF}} P(f_k) = \frac{1}{2}\sum_{f_k=f_{\min}}^{f_{\max}} P(f_k)$$

- **High-Frequency Power Ratio (HF$_{\text{ratio}}$):** The proportion of spectral power residing in the high-frequency band (150–400 Hz) relative to total power (15–500 Hz), capturing the recruitment patterns of fast-twitch (Type II) motor units:

$$\text{HF}_{\text{ratio}}^{(i)} = \frac{\int_{150}^{400} P(f)df}{\int_{10}^{500} P(f)df}$$

- **Dimitrov Fatigue Index (FI$_{\text{Dimitrov}}$):** The ratio of weighted spectral moments where the numerator weights the low frequency power and denominator weights the high frequency power. This index is highly sensitive to fatigue-induced spectral shifts[77]:

$$\text{FI}_{\text{Dimitrov}}^{(i)} = \frac{\int_{5}^{500} f^{-1} P(f)df}{\int_{5}^{500} f^{5} P(f)df}$$

**C. Discrete Wavelet Transform (DWT) Decomposition**

To preserve localized time-frequency energy distributions[78-80], sEMG signals were decomposed using a 5th-order Daubechies mother wavelet (db5) across 9 levels ($k \in \{1, \dots, 9\}$, spanning detail coefficients $D_1$ to $D_9$ from 2–1024 Hz). The total detail energy at level $k$ during trial $i$ was:

$$P_{\text{DWT},k}^{(i)} = \sum_{j} | d_{k,j}^{(i)} |^2$$

Linear regression slopes across the 15 trials were computed for each wavelet decomposition level to capture scale-specific temporal evolution.

**D. Non-Linear Signal Complexity: Sample Entropy (SampEn)**

Sample Entropy measures time-series complexity and regularity without assuming strict stationarity[81]. For a discrete sEMG sequence $\mathbf{x} = \{x_1, \dots, x_N\}$, embedding vectors of length $m$ are defined as $\mathbf{u}_m(i) = [x_i, \dots, x_{i+m-1}]$. The distance between vectors is given by the Chebyshev norm:

$$d[\mathbf{u}_m(i), \mathbf{u}_m(j)] = \max_{k \in [0, m-1]} | x_{i+k} - x_{j+k} |$$

Let $B^m(r)$ denote the probability that two sequences of length $m$ match within tolerance $r$, and $A^m(r)$ the probability for length $m + 1$. SampEn is defined as:

$$\text{SampEn}(m, r, N) = -\ln\left(\frac{A^m(r)}{B^m(r)}\right)$$

Parameters were fixed to $m = 2$ and $r = 0.2\sigma_x$ (where $\sigma_x$ is the standard deviation of the 20 s steady-state sEMG window).

E. **Hemodynamic Features** ($StO_2$)

- **Baseline Oxygenation (**$StO_{2,\mathrm{base}}$**):** Median tissue saturation index during the pre-exercise resting period.

- **Mid-Task Oxygenation Change (**$\Delta StO_{2,\mathrm{mid}}$**):** Relative change in mean clenching $StO_2$ during trials 4–6 compared to baseline (adapted from the pre-registered model where this feature was selected based on a data-driven feature-selection approach).

- **Hemodynamic Recovery Slope (**$\Delta StO_{2,\mathrm{slope}}$**):** Linear regression slope of the trial-by-trial difference between steady-state clenching and resting medians ($\Delta StO_2^{(i)} = StO_{2,\mathrm{clench}}^{(i)} - StO_{2,\mathrm{rest}}^{(i)}$), reflecting microvascular reperfusion efficiency.

**3. Baseline Model Configurations and Feature Sets**

Four distinct classical models were constructed to evaluate the specific predictive capacity of different feature domains (summarized in **Table S2**).

- **Model 1: Pre-Registered Partial Least Squares (PLS-Preregistered)**
  This model was developed using a portion of the Control-train dataset (n=70) with 10-fold cross-validation and sequential feature selection. The preregistration is available at https://osf.io/fwne7. This baseline model adheres to the preregistered protocol using 5 selected features:

$$\mathbf{x}_{\mathrm{PLS}} = \left[ |\,\mathrm{SampEn}_{15} - \mathrm{SampEn}_{1}\,|, \mathrm{sign}(\Delta StO_{2,\mathrm{slope}}), \mathrm{sign}(\mathrm{Slope}(P_{\mathrm{DWT},D1})), \mathrm{sign}(\mathrm{Slope}(\mathrm{SampEn})), \Delta StO_{2,\mathrm{mid}} \right]^T$$

  Predictions were generated using Partial Least Squares Canonical Regression (PLSCanonical with one component) following $Z$-score standardization. Convergence tolerance was tuned via nested 10-fold cross-validation (tol $\in \{10^{-5}, 10^{-6}, 10^{-7}, 10^{-8}\}$).

- **Model 2: Median Frequency Spectral Shift (Linear-Spectral Shift)**
  This model explicitly tests the classical hypothesis that myalgia manifests primarily as a progressive downward shift in median frequency[51,82]. It utilizes a 4-dimensional feature vector:

$$\mathbf{x}_{\mathrm{SpecShift}} = \left[\mathrm{MDF}_{\mathrm{slope}}, (\mathrm{MDF}_{15} - \mathrm{MDF}_1), \overline{MDF}, \mathrm{MDF}_{\mathrm{MVC}}\right]^T$$

  Predictions were generated via Ridge Regression, with the L2 regularization penalty ($\alpha \in [10^{-4}, 10^3]$) tuned via nested 10-fold cross-validation.

- **Model 3: Comprehensive Linear Regression (Linear-EMG)**
  A robust Ridge regressor incorporating the full 23-dimensional vector of handcrafted motor and metabolic features. Hyperparameters were tuned identically to Model 2.
- **Model 4: Gradient Boosted Decision Trees (XGBoost-EMG)**
  To evaluate non-linear decision boundary capabilities on the 23-dimensional feature space without utilizing deep sequence modelling, an XGBoost Regressor was trained (100 estimators, maximum tree depth = 3, learning rate = 0.05, subsample ratio = 0.8).

**4. Bilateral Modelling and Evaluation Protocol**

Independent estimators ($\mathcal{M}_{\text{Left}}$ and $\mathcal{M}_{\text{Right}}$), were trained to predict side-specific post-MVC subjective pain ratings. Models were evaluated across all benchmark settings detailed in the main text (Control-test hold-out, Control → mTMD zero-shot transfer, mTMD CV, mTMD → Control transfer, and all subjects CV). Predictive performance was quantified using Spearman's rank correlation ($\rho$), with statistical significance determined via non-parametric permutation testing (10,000 iterations).

### Table S2: Feature Matrix Summary by Model.

| Feature Category | Feature Description | PLS | Spec Shift | Linear | XGBoost |
|---|---|---|---|---|---|
| **Non-Linear Complexity** | Entropy Change Magnitude | X | | | |
| | Sign of SampEn Slope | X | | | |
| | Mean SampEn (across 15 trials) | | | X | X |
| | SampEn Slope (across 15 trials) | | | X | X |
| **Hemodynamics** ($StO_2$) | Sign of Clench-Rest $StO_2$ Difference Slope | X | | | |
| | Mid-Task $StO_2$ Change | X | | X | X |
| | Baseline $StO_2$ | | | X | X |
| | Clench-Rest $StO_2$ Difference Slope | | | X | X |
| **Wavelet (DWT db5)** | Sign of Level 1 Detail Power Slope | X | | | |
| | Detail Power Slopes Levels 1–9 (9 features) | | | X | X |
| **Spectral (PSD)** | Median Frequency (MDF) Slope | | X | X | X |
| | Net MDF Change | | X | | |
| | Mean MDF | | X | X | X |
| | MVC MDF | | X | | |
| | Mean Frequency (MNF) | | | X | X |
| | High-Frequency Power Ratio | | | X | X |
| | Dimitrov Fatigue Index ($FI_{Dimitrov}$) | | | X | X |
| **Time-Domain (Amplitude)** | Mean RMS (across 15 trials) | | | X | X |
| | RMS Slope (across 15 trials) | | | X | X |
| | MVC RMS | | | X | X |
| | Mean Zero-Crossing Rate (ZCR) | | | X | X |
| **Total Features (Per Side)** | | **5** | **4** | **23** | **23** |

### Supplementary Note 2: Prospective Validation Plan

Although we validated LENS in independent held-out datasets of healthy controls and individuals with chronic mTMD pain, it has not been validated in an entirely external dataset. To separate site, operator and hardware effects from biology, the frozen model should first be applied without retraining to a masseter cohort (n ≈ 50; 25 healthy controls and 25 participants with mTMD) recorded at an independent centre using an EMG system from a different manufacturer, with the primary endpoint pre-specified as the Spearman correlation between predicted and reported post-exertion pain and the model weights time-stamped before data collection begins. To test whether the signature is craniofacial-specific, the same protocol should be adapted to a limb or axial muscle in which graded fatiguing contraction can be standardized—for example the upper trapezius or the quadriceps—in healthy volunteers and in a matched chronic MSK pain cohort. To establish whether the calibration step is clinically practical, participants should be recorded on two occasions at least 24 h apart, permitting test–retest reliability of both the predictions and the adapted encoder weights to be quantified. To enable independent replication in advance of these studies, the frozen model weights, the preprocessing and evaluation pipeline, and anonymized data from the 152 participants who consented to sharing are publicly released.

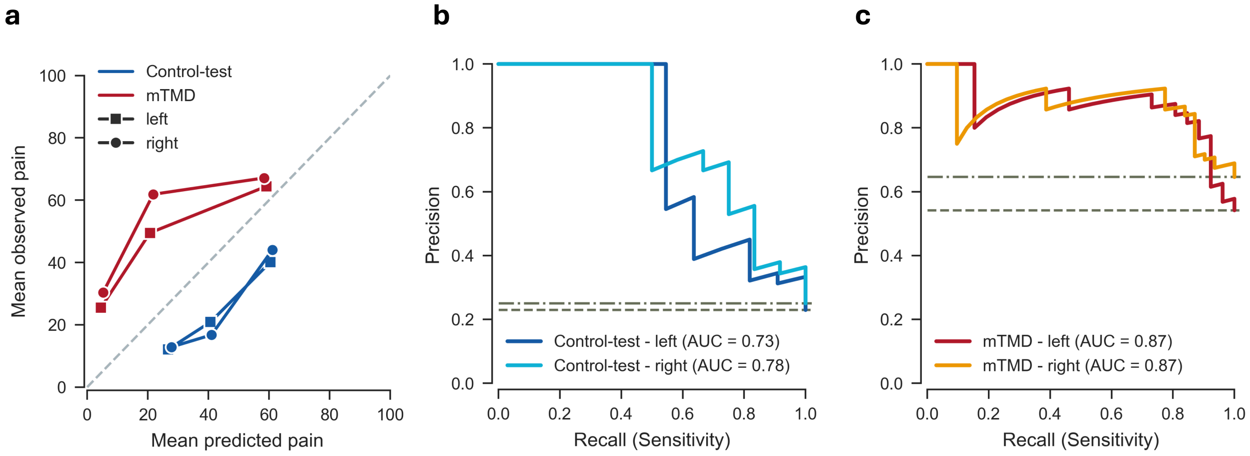


**Figure S1: Clinical calibration and precision-recall characteristics.**

**a.** Tercile-based calibration plot comparing mean predicted versus mean observed pain. The dashed diagonal represents perfect ideal calibration. The model exhibits a slight over-prediction in healthy controls (blue, below ideal) and an under-prediction of pain in mTMD (red, above ideal), while preserving monotonic rank ordering. **b-c.** Precision-Recall (PR) curves for the Control-test (**b**) and mTMD (**c**) cohorts at a VAS≥45/100 threshold. Horizontal dashed lines represent the no-skill baseline (dataset prevalence of positive samples: Control-test - left = 0.23, right = 0.25; mTMD – left = 0.54, right = 0.65).

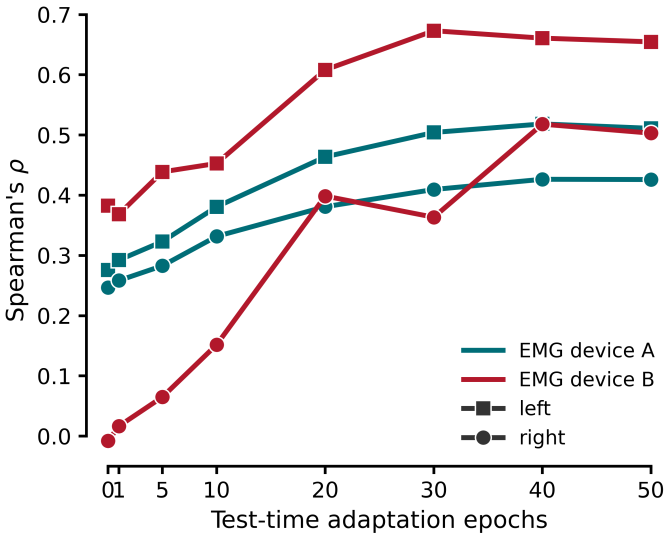


**Figure S2:** Performance (Spearman's ρ) as a function of TTA epochs for mTMD participants, stratified by recording device. The adaptation trajectory is consistent for both Device A (n=35) and Device B (n=19), confirming that TTA corrects for biological pathology rather than hardware artefacts.

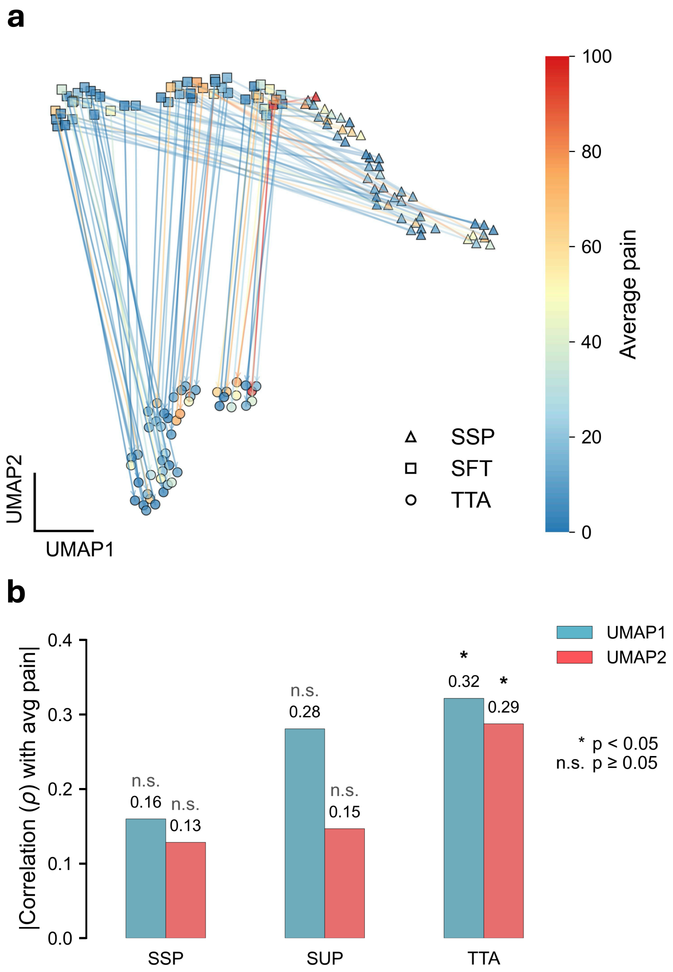


**Figure S3: Evolution of encoder embeddings across training stages**

**a,** UMAP projection of mean-pooled encoder embeddings across the three training stages (SSP, SFT, TTA). **b,** Correlation between the principal UMAP components and pain ratings. The structural alignment of the embeddings with the pain gradient improves progressively, peaking after Test-Time Adaptation.

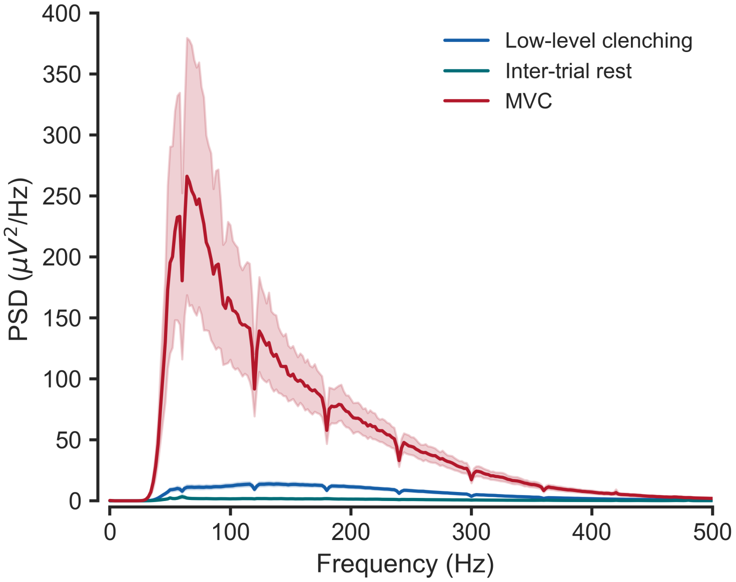


**Figure S4:** Power Spectral Density (PSD) of the raw EMG signal from the last 7 trials (trials used as input to the model) as well as the MVC period in healthy participants. EMG signal was bandpass filtered (40-850 Hz) as part of the pre-processing pipeline, and we used notch filters to attenuate the line noise at 60 Hz and its harmonics for visualization purposes. Inter-trial rest and MVC show higher power in lower frequencies (<100 Hz) whereas low-level clenching has the highest power in the 100-200 Hz range.

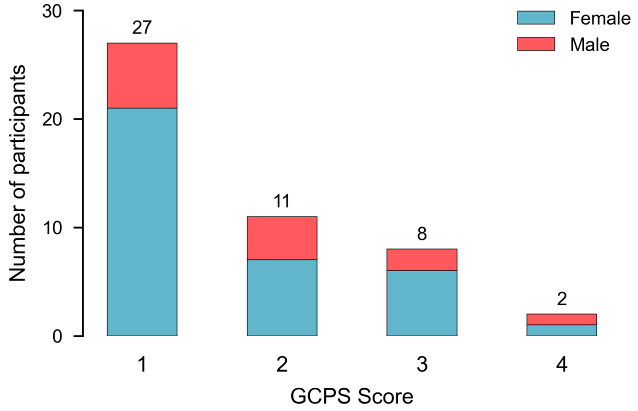


**Figure S5: Distribution of Graded Chronic Pain Scale (GCPS) in individuals with chronic mTMD.**

Bar plot illustrating the frequency distribution of pain severity and disability among participants in the mTMD cohort (n=54), assessed via the Graded Chronic Pain Scale. Participants are stratified into grades reflecting the spectrum of chronic pain severity: grade 0 (pain-free), grade 1 (low intensity pain, none-low pain-related disability), grade 2 (high intensity pain, none-low pain-related disability), grade 3 (moderately limiting), and grade 4 (severely limiting). This highlights the clinical heterogeneity of the chronic mTMD cohort evaluated in this study.

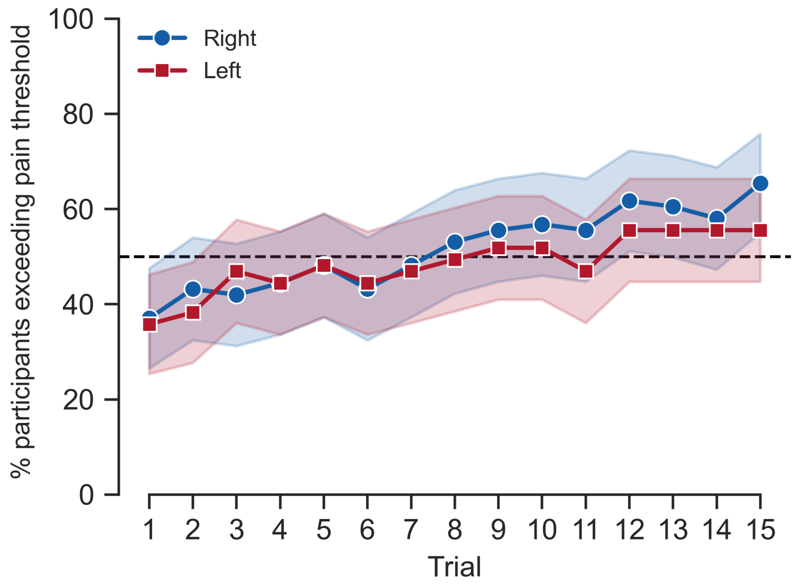


**Figure S6: Selection of training window based on pain onset.**

Proportion of participants in the Control-train cohort reporting a pain intensity exceeding the minimum perception threshold (5/100 VAS) across the 15 clenching trials. By trial 9 (dashed line), 50% of the participants exceed this threshold on both the left and right sides. Consequently, trials 9–15 were selected as the input window for the predictive model to ensure the network was trained on segments containing active nociceptive information.

**Table S3: Linear Mixed-Effects Models (LMM) evaluating the temporal decline of relative 150–400 Hz power.**

| Predictor | Coefficient (β) | Bootstrap CI | P-value |
|---|---|---|---|
| ***Model A: Pain-Driven Interaction*** | | | |
| Intercept | -2.22 | [-2.41, -2.04] | <0.001 |
| Trial (scaled 0–1) | -0.07 | [-0.25, 0.12] | 0.617 |
| Pain (scaled 0–1) | -0.28 | [-0.79, 0.20] | 0.209 |
| Trial × Pain | -0.68 | [-1.39, -0.07] | 0.025 |
| ***Model B: Group-Driven Interaction*** | | | |
| Intercept | -2.23 | [-2.3, -2.11] | <0.001 |
| Trial (scaled 0–1) | -0.25 | [-0.50, -0.06] | 0.026 |
| Group (mTMD) | -0.33 | [-0.73, 0.02] | 0.028 |
| Trial × Group | -0.23 | [-0.63, 0.15] | 0.252 |

Model A evaluates the continuous effect of subjective pain intensity (scaled 0-1) on the rate of high-frequency power decline over 15 clenching trials (scaled 0-1). Model B evaluates the categorical effect of pathological diagnosis (mTMD vs. control) on the same temporal decline. Both models include random intercepts and random slopes for the trial variable grouped by subject. Interaction p-values indicate that spectral decline is driven by post-MVC pain rather than pathological group status. 95% Confidence Intervals were generated via 5,000 subject-level block bootstrap iterations.

**Table S4. Relationships among reported pain, reported fatigue and LENS predictions.**

| Cohort | Side | $\rho(\hat{y}, y)$ | $\rho(y, fatigue)$ | $\rho(\hat{y}, fatigue)$ | $\rho(\hat{y}, y \mid fatigue)$ | $\rho(\hat{y}, fatigue \mid y)$ |
|---|---|---|---|---|---|---|
| **Control-test** | Left | **0.52** *(p=0.0001)* | 0.24 *(p=0.05)* | 0.18 *(p=0.1)* | **0.50** *(p=0.0003)* | 0.07 *(p=0.63)* |
| | Right | **0.48** *(p=00005)* | 0.17 *(p=0.12)* | 0.10 *(p=0.24)* | **0.48** *(p=0.0007)* | 0.02 *(p=0.87)* |
| **mTMD** | Left | **0.61** *(p=0.0001)* | **0.79** *(p=0.0001)* | **0.54** *(p=0.0001)* | **0.37** *(p=0.01)* | 0.14 *(p=0.35)* |
| | Right | **0.48** *(p=0.0005)* | **0.81** *(p=0.0001)* | **0.51** *(p=0.0002)* | 0.15 *(p=0.33)* | 0.23 *(p=0.11)* |

Spearman rank correlations (ρ) with p-values for each cohort and masseter, together with partial correlations of predictions with pain controlling for fatigue, and of predictions with fatigue controlling for pain. In the Control-test cohort, pain and fatigue are largely separable ($\rho^2 < 0.06$), and partial correlations are interpretable. In the mTMD cohort the two reports are near-collinear ($\rho^2 = 0.63$–$0.66$), so partialling fatigue removes the majority of the rank variance in the pain label and the resulting partial coefficients should not be read as measures of specificity. y: post-MVC pain intensity; $\hat{y}$: LENS predictions.
